# A fresh look at an old concept: Home-range estimation in a tidy world

**DOI:** 10.1101/2020.08.19.256859

**Authors:** Johannes Signer, John Fieberg

## Abstract

1. A rich set of statistical techniques have been developed over the last several decades to estimate the spatial extent of animal home ranges from telemetry data, and new methods to estimate home ranges continue to be developed.
2. Here we investigate home-range estimation from a computational point of view and aim to provide a general framework for computing home ranges, independent of specific estimators.
3. We show how such a workflow can help make home-range estimation easier and more intuitive, and we provide a series of examples illustrating how different estimators can be compared easily, so that one can perform a sensitivity analysis to determine the degree to which the choice of estimator influences qualitative and quantitative conclusions.
4. By providing a standardized, tidy implementation of home-range estimators, we hope to equip analysts with the tools needed to explore how estimator choice influences answers to biologically meaningful questions.

## Introduction

The biological concept of an animal’s *home range* has served as a useful construct for organizing our thinking about how animals use and interact with space since the time of Darwin (Kie et al. 2010; Horne et al. 2020). Today, most people associate the term home range with Burt (1943)’s definition, “that area traversed by the individual in its normal activities of food gathering, mating and caring for young. Occasional sallies outside the area, perhaps exploratory in nature, should not be considered as in part of the home range.” A variety of statistical and modeling approaches have been developed to quantify the spatial extent and intensity of landscape use by individual animals and to gain insights into factors that structure their home ranges (see e.g., Powell 2012 and associated papers in a special feature on the topic). Recently, Fleming et al. (2016) and Horne et al. (2020) have argued for classifying statistical home-range methods by whether they estimate one of two estimation targets: the *range distribution* or long-term (equilibrium) distribution that would result from an animal continuing to move in a consistent manner and an *occurrence distribution* that captures the path of movement an animal takes during a specific observation window, along with its uncertainty. This dichotomy is appealing from a theoretical point of view, and several new statistical estimators have been developed for targeting these quantities while also addressing issues related to autocorrelation, a prominent feature of modern day Global Positioning System (GPS) data (Fleming et al. 2014, 2015, 2016).

Despite these advances, many biologists continue to use a variety of “old” estimators (e.g., minimum convex polygons [MCP]; Mohr 1947; or kernel density estimators [KDE] that assume independent location data; Worton 1989) without explicit discussion of a particular estimation target (e.g., Froy et al. 2018; Ranc et al. 2020). We suspect there may be multiple reasons why, including: 1) some ecologists may not be familiar with recent literature on home-range estimators; 2) current estimators that account for autocorrelation require an extra step of fitting an animal movement model to location data, which can also can take considerable time and computational resources when applied to large data sets involving many animals; some researchers may feel this extra step is unnecessary or they may not feel confident in their use of these methods; and 3) researchers may be interested in estimating something other than a range or occurrence distribution. The new methods developed by Fleming and co-authors are major contributions to this area of research, and these authors have done a nice job providing open-source software and training for implementing their estimators (Fleming and Calabrese 2020; Calabrese et al. 2020). We conjecture, however, that many biologists continue to view traditional home-range estimators as convenient, though imperfect, indices that capture the spatial extent of the area used by individuals during specific tracking periods. For convenience, and as is common in the literature, we will refer to the suite of methods used in this context as *home-range estimators*, even though these methods may have different statistical estimation targets (Horne et al. 2020).

Whereas there are many studies that compare different methods for quantifying space use with the goal of determining a single “best” estimator (e.g., Lichti and Swihart 2011; Walter, Onorato, and Fischer 2015; Noonan et al. 2019), we aim in this paper to be largely estimator agnostic. As we have argued previously, we think researchers should carefully consider estimators and their properties (e.g., variance or statistical power), and choose one or more depending on the specific biological questions of interest (Fieberg and Börger 2012; Signer et al. 2015). We also encourage users to consider multiple estimators, when possible, to evaluate the sensitivity of their results to estimator choice.

To accomplish this goal, we propose a general and consistent framework for home-range estimation that should be able to accommodate *most* home-range estimators. We propose two classes of home-range estimators and a set of properties for each class. Having a standardized treatment of home-range estimators facilitates their computation, visualization, and comparisons among estimators. This proposal goes hand in hand with calls for more reproducible and standardized workflows in (wildlife) ecology (Gula and Theuerkauf 2013; Lewis, Vander Wal, and Fifield 2018; Archmiller et al. 2020). After introducing the framework conceptually, we demonstrate how to estimate home-ranges using this framework following the principles of tidy data (Wickham and others 2014) using the R package `amt` (Signer, Fieberg, and Avgar 2019; R Core Team 2020) and a previously published data set of fishers from New York (LaPoint et al. 2013a).

### A conceptual framework for home ranges

Home-range estimators can be divided into two classes: geometric and probabilistic estimators (Figure 1; Fleming et al. 2015). Geometric estimators are constructed following a set of rules and are often hull-based, i.e., the home range is a polygon that is constructed using (all) points where an animal was observed. Typical examples of geometric estimators are minimum convex polygons (Mohr 1947) or local convex hulls (LoCoH; Getz and Wilmers 2004). On the other hand, probabilistic estimators have an underlying probabilistic model and estimate an utilization distribution, the two-dimensional relative frequency distribution of an animal’s spatial locations (Van Winkle 1975), from which a hull-based home range can be retrieved for a given isopleth level. Typical examples of probabilistic home-range estimators include uniform or bi-variate normal models (Van Winkle 1975; Horne and Garton 2006), traditional KDEs (Worton 1989; Fieberg 2007), and autocorrelated KDEs (aKDE; Fleming et al. 2015).

**Figure 1:**
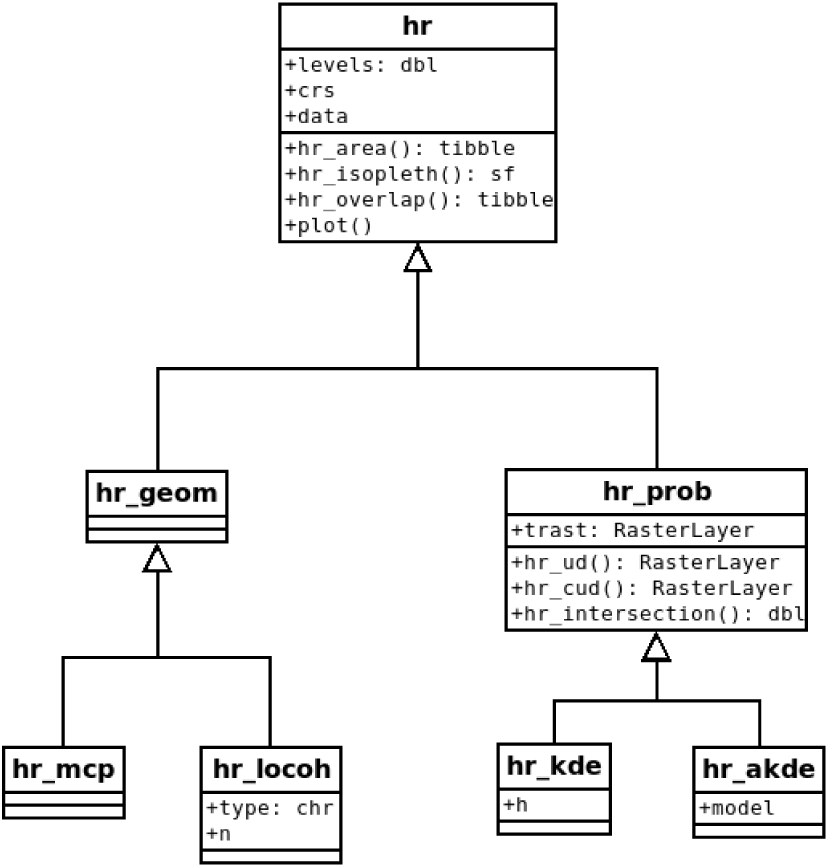
Proposed class diagram for home-range estimators. All home-range estimators will have common attributes (`levels`, `crs` and `data`) and common global methods (`hr_area()`, `hr_isopleth()`, `hr_overlap()`, and `plot()`). Probabilistic estimators will also have a common attribute, `trast`, and several additional common methods (`hr_ud()`, `hr_cud()`, and `hr_intersection()`). Lastly, each individual estimator can have additional properties or methods (e.g., `model` for `hr_akde()`).

Each home-range estimator, regardless of its class, has several attributes (values stored within the object) and methods (functions to work with the estimate). Estimators of both classes should have the following three attributes: the coordinate reference system (`crs`), the data that were used to construct the home-range estimate (`data`), and the home-range isopleths or levels (`levels`). The coordinate reference system is inherited from the data used to estimate the home range and will be needed to ensure that the home range is correctly positioned in space and that the units of the home-range area are correct. The attribute `data` contains the original data used to calculate the home range which can be especially useful for plotting home ranges. Finally, home-range areas are calculated for a pre-specified home-range level (or isopleth). For probabilistic estimators, the cumulative distribution function of the utilization distribution is truncated at given quantile, with associated 1 *− α* level. For hull-based methods the outermost points are excluded (given it is possible to identify the outermost points). It is common to use the 95% isopleth to determine the home range (although arguments for 90% levels exist e.g., Börger, Franconi, Ferretti, et al. 2006a). Further, we propose the following methods for all home-range estimators: `hr_area()` to calculate the home-range area, `hr_isopleth()` to calculate the home-range isopleths at the specified `levels`, `hr_overlap()` to calculate the home-range overlap between two or more home ranges, and `plot` to plot the home range.

In addition to the four global methods, specific methods or classes can have additional properties and/or methods. Probabilistic home ranges, for example, have a property that gives information about the spatial extent and resolution of the utilization distribution (here and in `amt` this is termed template raster, or `trast` for the argument name). Probabilistic home ranges should also have a method to obtain the utilization distribution (`hr_ud()`), the cumulative utilization distribution (`hr_cud()`), and to quantify volumetric intersections of two utilization distributions (`hr_intersection()`). Examples of estimator-specific properties are the number of neighbors used for the local convex hull method, the bandwidth for kernel density estimation, or the movement model used for autocorrelated kernel density estimation (Fig. 1).

An established set of methods makes it easy to work with home ranges. For example, the function `hr_area()` in the package `amt`, will always return a `tibble` (Müller and Wickham 2020) with two columns (the home-range level and area), regardless of the estimator. A `tibble` is very similar to a `data.frame` in R (i.e., a two-dimensional data structure) but with improved properties. A `tibble` makes it is easy to work with list-columns, which as we will demonstrate later in this paper, help to facilitate analyses of data from multiple animals or sampling instances. Similarly, the function `hr_isopleth()` in `amt` will always return a `tibble` with a simple feature column of class `sfc_POLYGON` from the `sf` package (Pebesma 2018), allowing further GIS-related work.

### One individual or sampling instance

In the first set of examples, we demonstrate how home ranges and derived quantities can be calculated for a single individual or sampling instance (e.g., a home range for one individual using data collected during a single tracking period). We use a data set containing locations of fisher from New York, USA (LaPoint et al. 2013a). These data are freely available from Movebank (LaPoint et al. 2013b), and include observations of six individuals (three males and three females) tracked between January and March 2011, with a sampling rate of 10 minutes or less. We use a preprocessed data set here. All steps to prepare the data set are provided in Supplement 1. For the first few examples, we will use data from one female (F1); for the second set of examples, we will use data from all six individuals.

First we load required packages, including `amt` for calculating home ranges (Signer, Fieberg, and Avgar 2019), `tidyverse` for data manipulation (this includes `ggplot2` for plotting; Wickham 2016, 2017), and `lubridate` for working with dates (Grolemund and Wickham 2011). After loading the data, we use the function `make_track()` to create a `track` – an object class used by `amt`, and then use `filter` to filter only those relocations that belong to the fisher where `id == “F1”`.

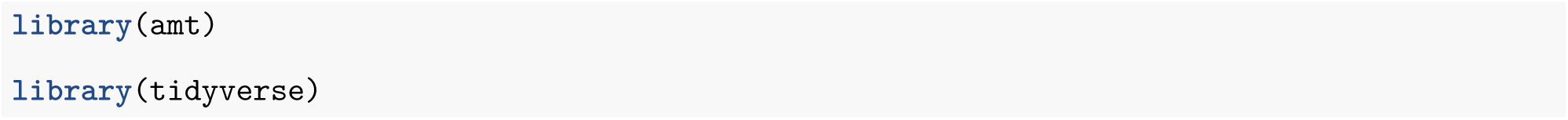

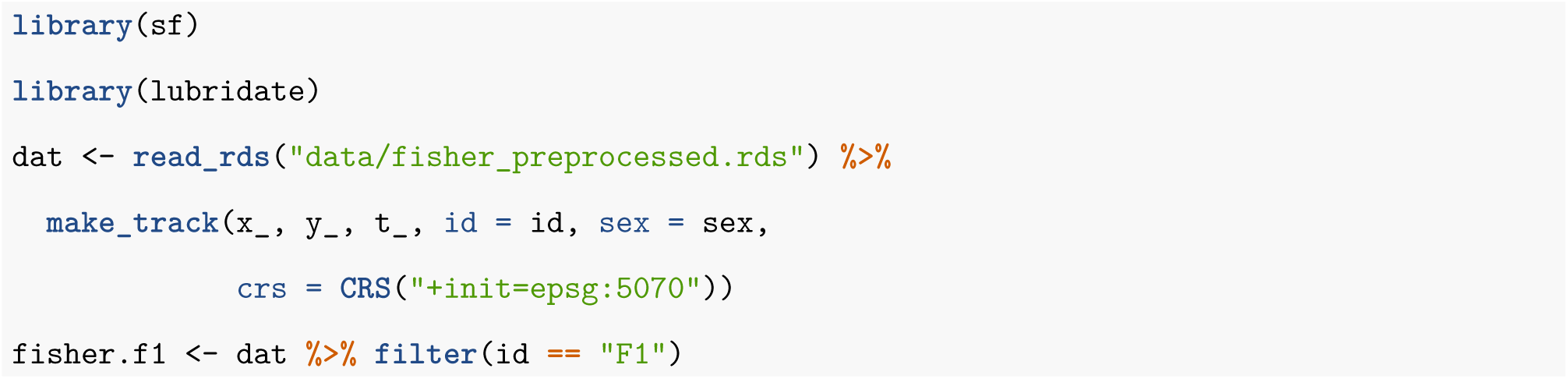

With the `fisher.f1` data set, we can now calculate different home-ranges estimates. We demonstrate by calculating MCP and KDE home ranges here (with the default reference bandwidth). For both home-range estimators, we estimate home ranges at two different home-range levels (50% and 95%).

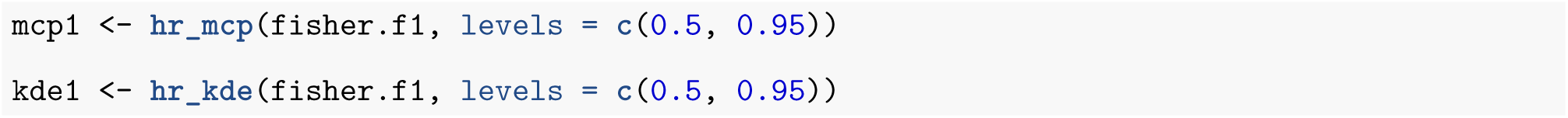

Results from applying any estimator in `amt` – not just the ones illustrated here – are stored in a `list`. All estimators have three entries in common: `crs`, `data` and `levels`; `crs` stores the coordinate reference system of the home range estimate, inherited from the data used to estimate the home range. The attribute `data` contains the track that was used to estimate the home range (a `track_xy*` from package `amt`), unless during estimation the argument `keep.data` was set to `FALSE`, then the attribute `data` is `NULL`. Finally, the argument `levels` contains the home-range levels that were used when estimating the home range.

All estimators also have at least four generic functions for working with the results and for basic plotting. The `plot()` function plots the home range isopleths with the observed points unless `keep.data` is set to `FALSE` or the argument `add.points` is set to `FALSE`. Below, we plot KDE- and MCP-based home ranges. When plotting the MCP, we use the arguments `add.relocations = FALSE` to avoid plotting the observed locations twice; further, we set the argument `add = TRUE` to draw the MCP home range to the existing plot and `border = “red”` to distinguish the KDE home range from the MCP home range by border color (Fig. 2).

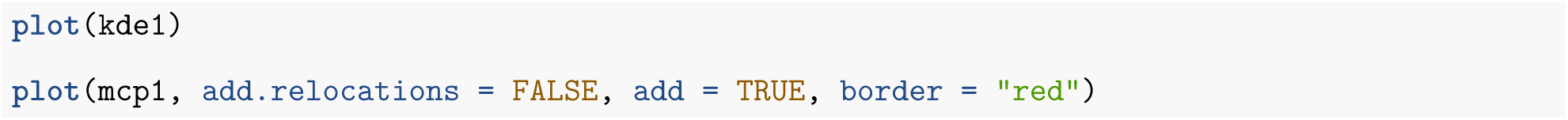

**Figure 2:**
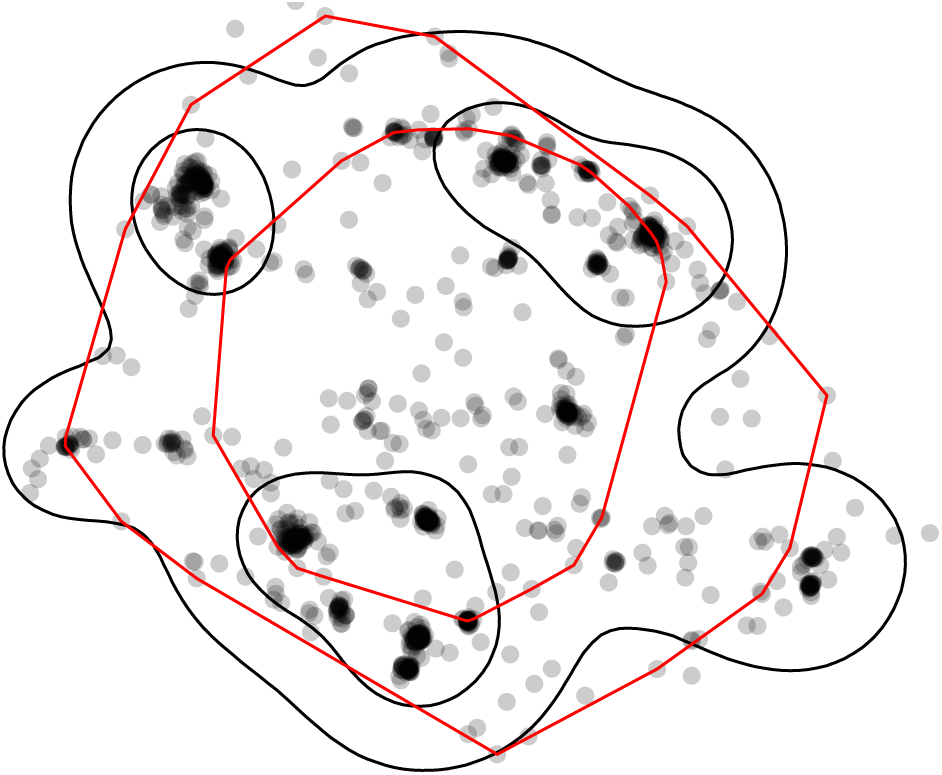
Points were the animal was tracked (black points) overlaid with kernel density (black lines) and minimum convex polygon (red lines) home ranges at two levels (50% and 95%). Note, that whereas home ranges delineated using the 95% level are relatively similar, the home ranges at the 50% level are very different.

Furthermore, we can now continue to work with these home-range estimates. For example, we can query the home-range area with the function `hr_area()`, which returns a `tibble` with two columns: the home-range level and the corresponding area.

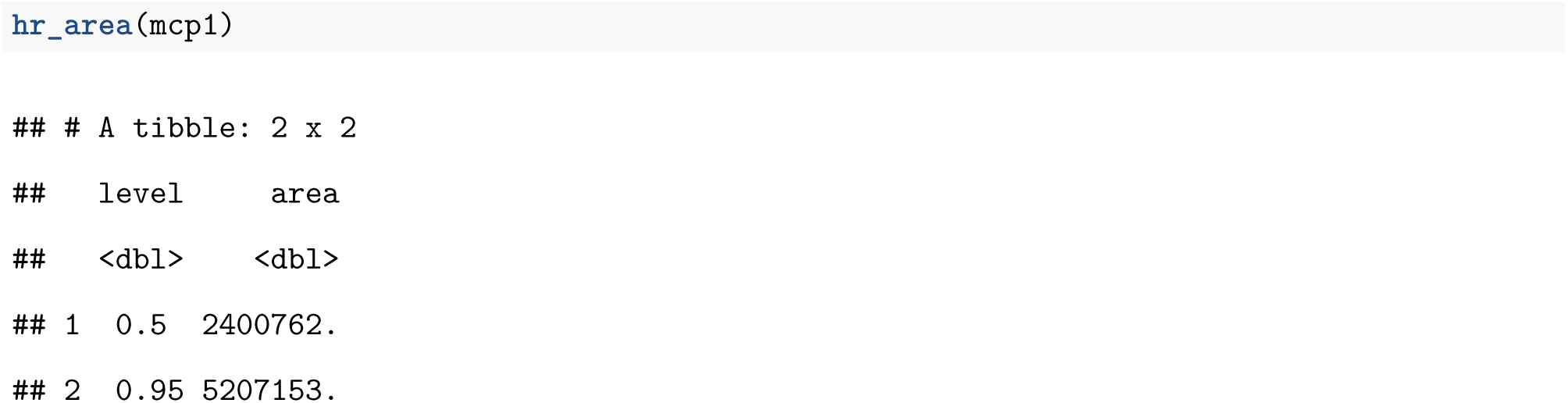

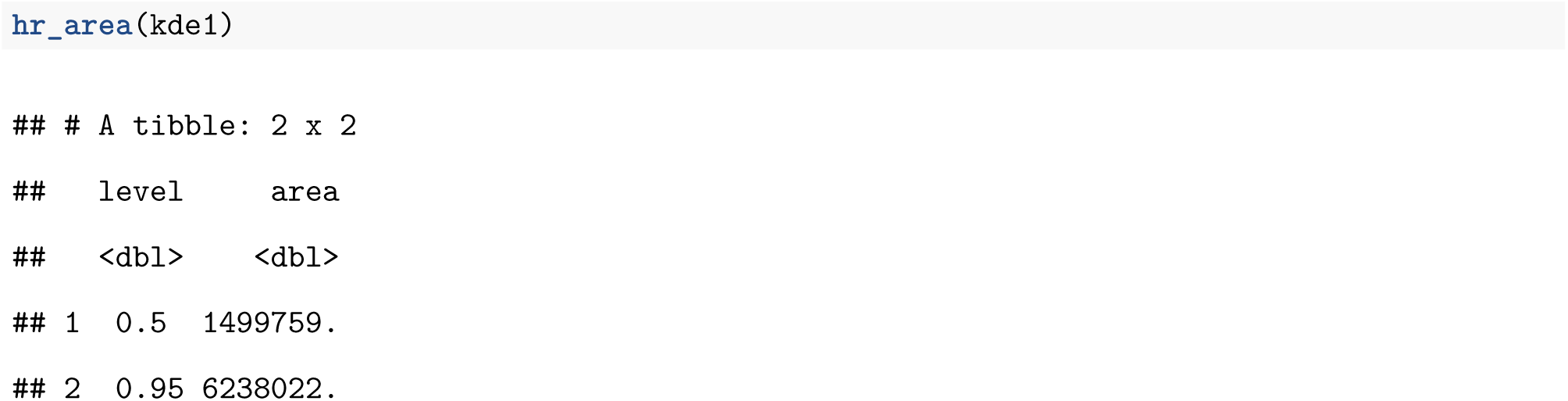

The function `hr_isopleth()` returns a `tibble` with a simple feature column of class `sfc_POLYGON` from the `sf` package (Pebesma 2018), which we can use to conduct further spatial analyses, visually inspect the home range, or export it to a GIS.

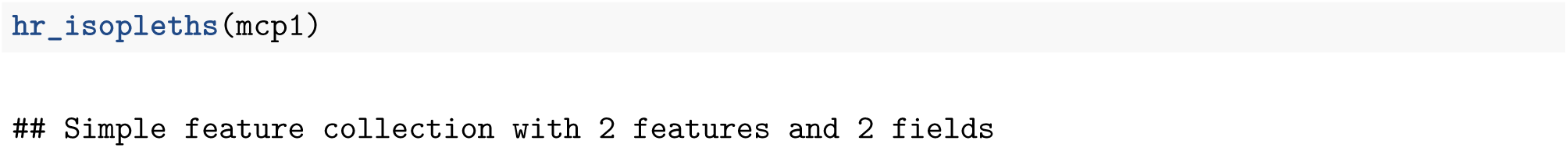

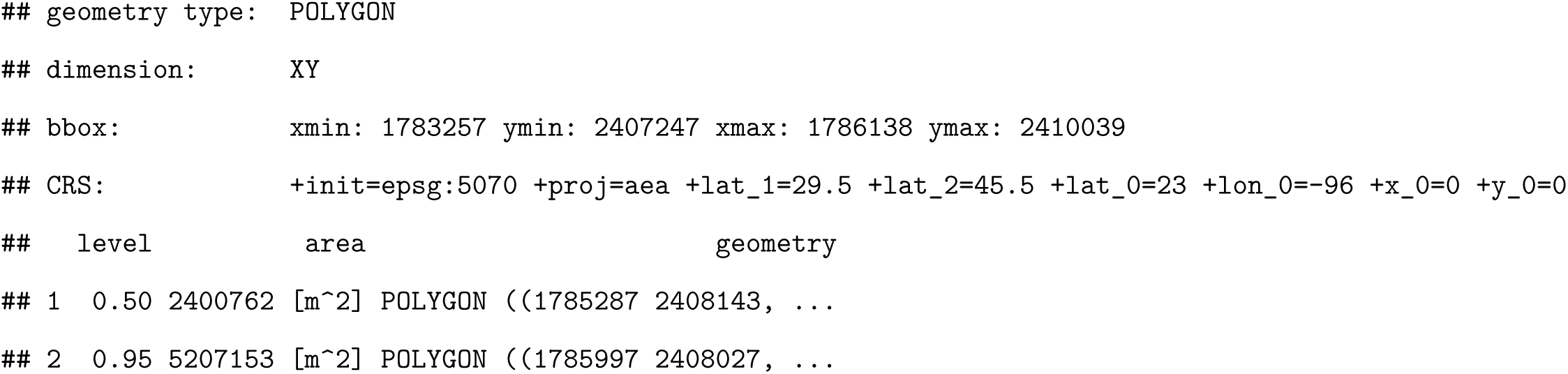

Finally, we may want to calculate the extent of overlap between two or more home ranges (Fieberg and Kochanny 2005), for which we provide the function `hr_overlap()`. This function can be used to calculate overlap between any two sampling instances (e.g., time periods, animals or even estimators). Below we calculate overlap of the MCP and KDE home ranges, which were previously estimated.

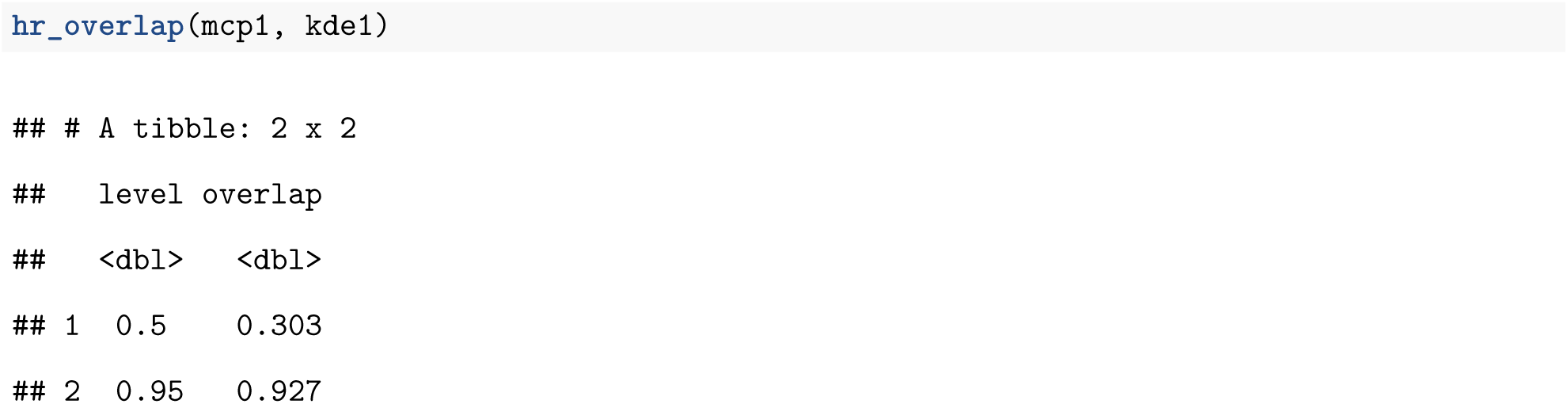

`hr_overlap()` always calculates the fraction of the first home range (i.e., the first argument) that is intersected by the second home range (second argument). Hence, changing the order of arguments will lead to a different result (Fieberg and Kochanny 2005).

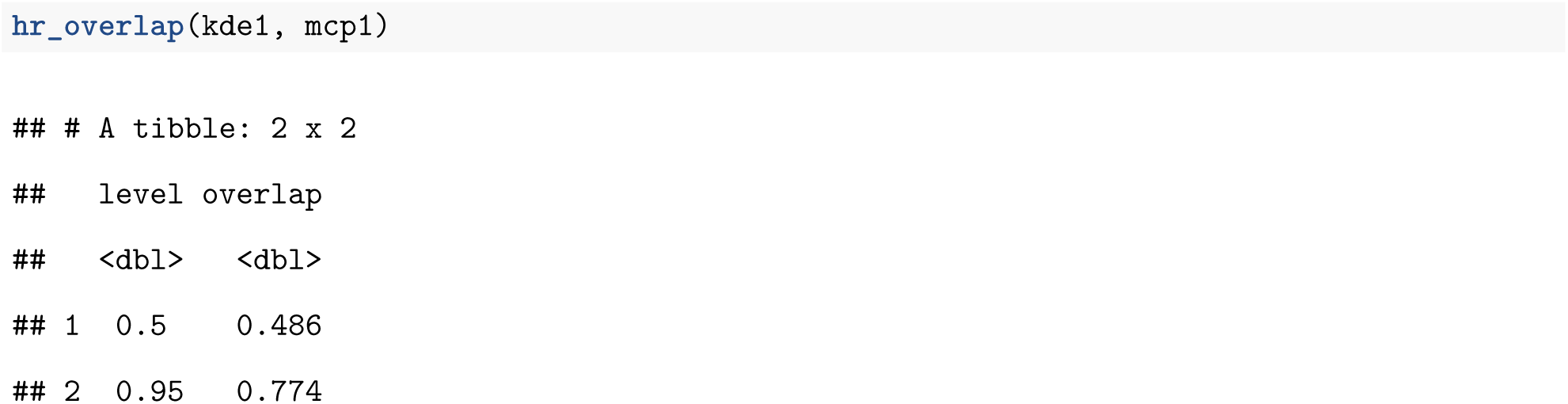

For probabilistic estimators, several other home-range overlap indices have been proposed (Fieberg and Kochanny 2005); these are also implemented in the `amt` package.

### Many individuals or sampling instances

Most telemetry studies collect data on several individuals and/or during several sampling instances (e.g., time periods). To compare estimates across sampling instances and to facilitate population-level inference, it is therefore important that the methods discussed so far for individual home ranges scale easily to situations with many animals and/or time intervals. The package `amt` does not provide an infrastructure for multiple instances, but instead relies on general data structures for such situations: *list columns* from the `tibble` package. A list column is a column of a `tibble` that contains a list. And a list, in turn, is a very flexible data structure that can hold almost any object. Thus, we can store results from applying home-range estimators in a list column of a `tibble`, together with meta-information (such as the name of the individual, its sex or age, or the time period it when was tracked) in other columns.

To demonstrate list columns, we will consider the full data set containing locations from all six fishers and illustrate workflows addressing three different example questions:

1. Do estimates of home-range size differ between sexes?
2. Is there a correlation between environmental covariates and estimates of home-range size?
3. How do daily “home ranges” change over time?

The aim of these examples is twofold: 1) we illustrate the benefit of standardized classes for home-range estimation (Fig. 1) and list columns, and 2) we highlight that some results are sensitive to the choice of estimator whereas others are not. In particular, estimates of home-range size tend to vary considerably among estimators, but relative comparisons over time or space are often robust to estimator choice (Signer et al. 2015).

All three of the above questions require that we iterate over several animals (question 1 and 2), and animals and days (question 3). List columns that organize data for each individual or sampling instance provide a simple way to facilitate these analyses. The function `nest()` from package `tidyr` (Wickham and Henry 2020) can be used to create a list column; `nest()` only requires the name of the list column and the columns that should be nested into the list. When using the syntax `nest(data = c(x_, y_, t_))`, below, all columns that are not named in the `nest()` call act as grouping variables. In our first example, the only column not listed is `id`, so it serves as a grouping variable; later we will show to group by more than one grouping variable. Alternatively, we could have used `nest(data = -c(id))` to specify that we want to use `id` as a grouping column, and that all other columns should be nested. These two approaches will result in identical results. Also note that we could choose a different name for the list column (i.e., it does not need to be labeled `data`).

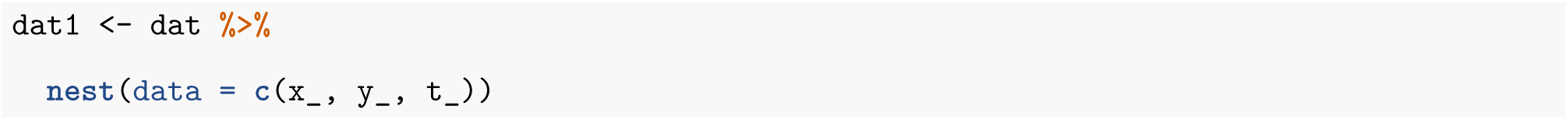

The result of `nest` is a `tibble` with three columns: `id`, `sex` and `data`.

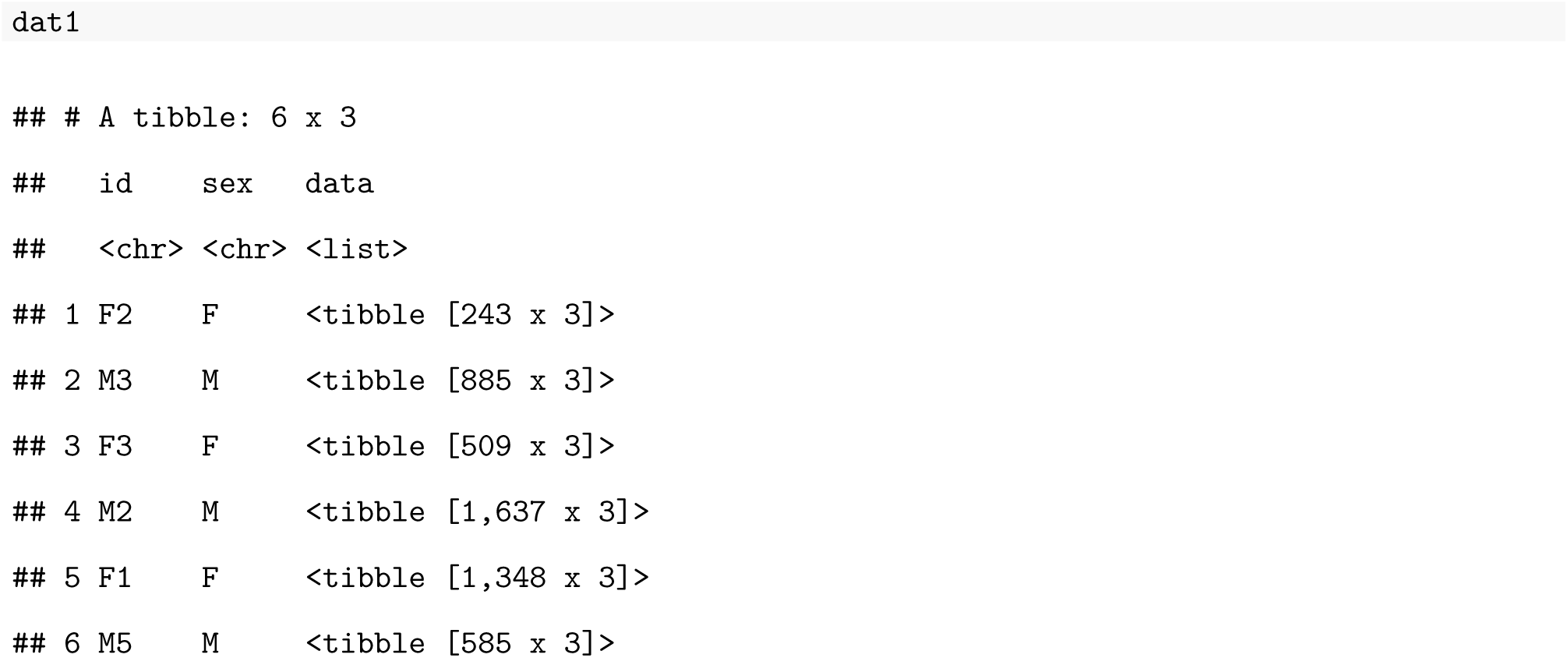

`data` is a list column that contains a tibble with all the relocations for a given animal (`id`). This list can be accessed in the regular way. For example to obtain relocations for the first animal, we can use:

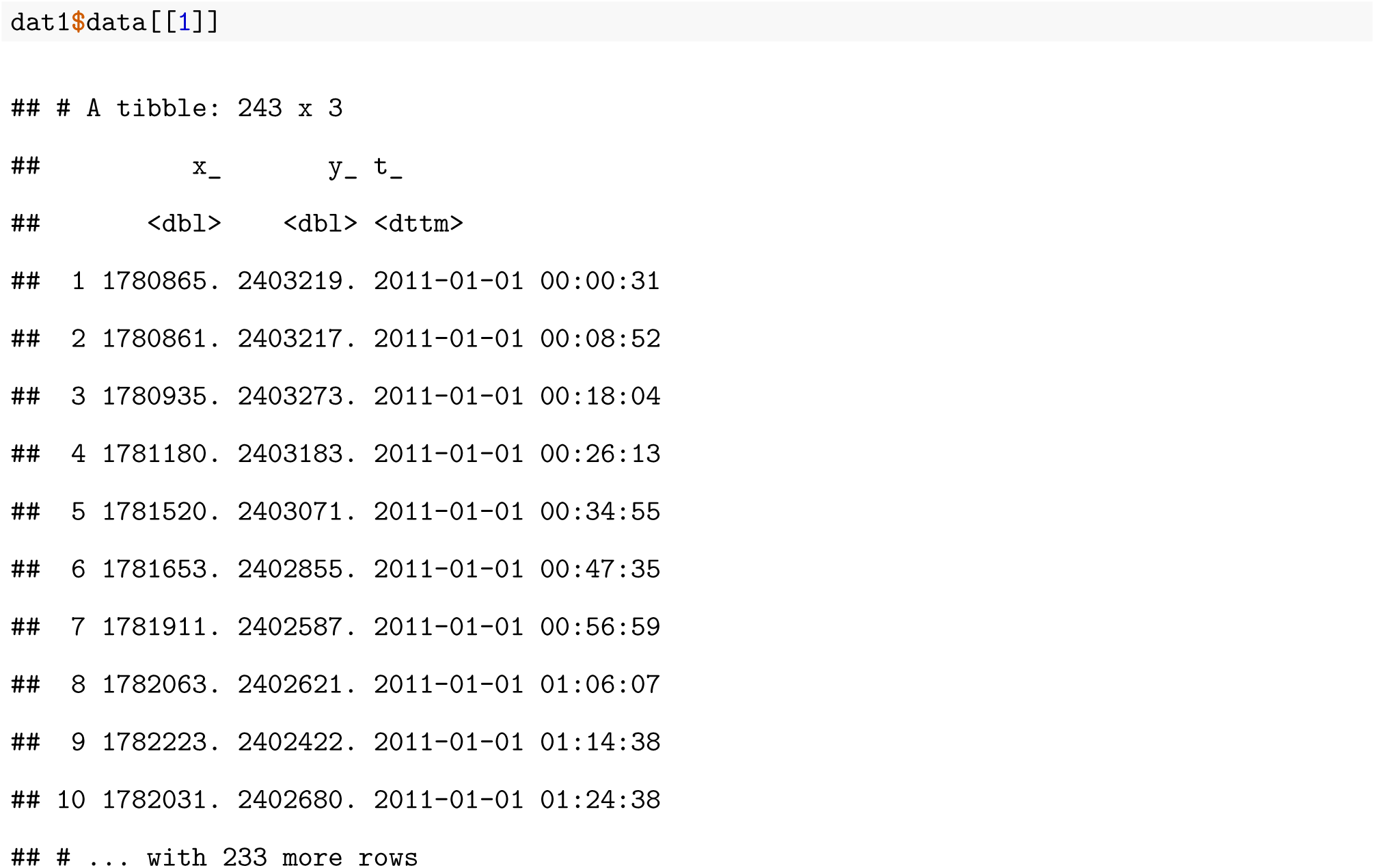

With the `mutate()` function, we can create a new list column that contains the home-range estimates for each animal. To achieve this goal, we have to iterate over each element in the column `data`, apply a home-range estimator, and save the result in a list. In base R, the function `lapply()` is well suited to this task. An alternative is the function `map()` from the `purrr` package (Henry and Wickham 2020).

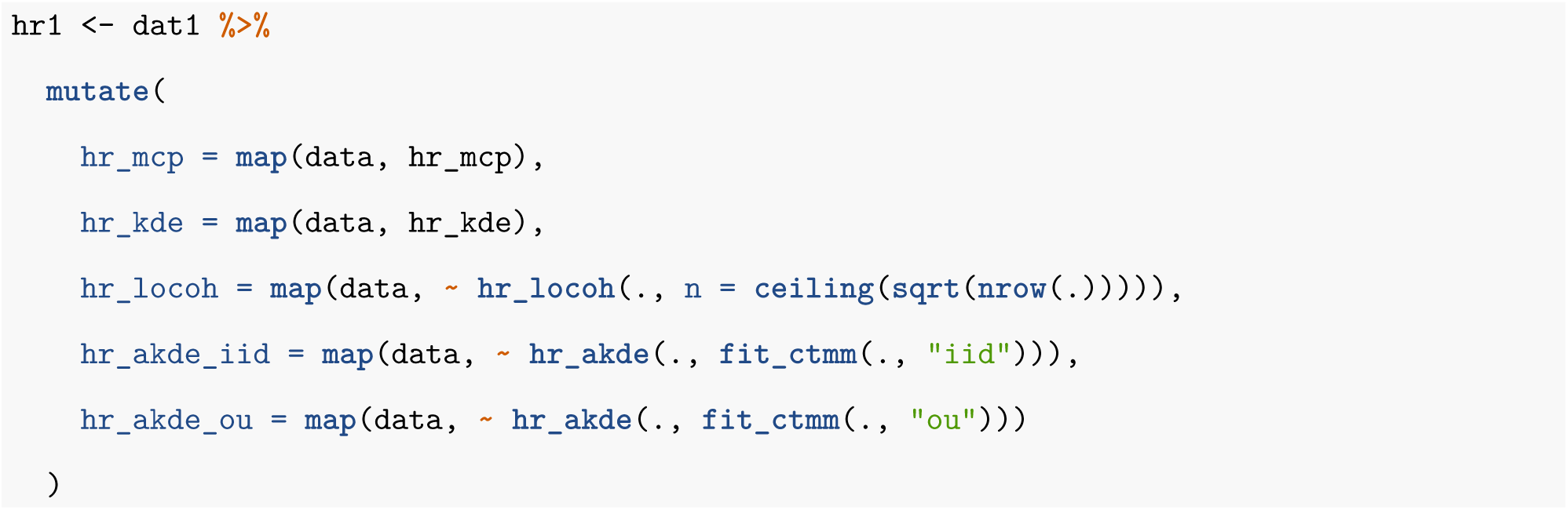

The function `map()` always iterates over a data structure (e.g., a vector or a list) that is provided as its first argument. The second argument to `map()` is a function that is to be applied to each element of this data structure. There are three different snytaxes we may use to specify this function: 1) we can simply supply the function name, as was done for the new column `hr_mcp`. In this case, `map()` is given the tracking data of each animal (stored in the column `data`) and the data set for each animal is then passed to the function `hr_mcp()`. This syntax works because the function `hr_mcp()` does not require the specification of further arguments (note, the default value of 0.95 for the home-range level is used). 2) A formula (`∼`) notation can be used to pass a function to `map()`. The advantage of this notation is that it is possible to access the data under evaluation – i.e., the relocation data of the current animal can be accessed either through a ., as we illustrate above, or through the predefined variable `.x` or `..1`. For example, for the local convex hull method, we want to choose `n` (the number of neighbors) as the square root of the number of observations. Thus, we count the number of rows with `nrow(.)` and then take the square root. Similarly, for the aKDE home-range estimator, we first want to fit a continuous-time movement model to the relocation data and then use this model when estimating the home range. Thus, we first pass the data, again using the ., to the function `fit_ctmm()` and then pass the result to the function `hr_akde()`. 3) `map()` can be used analogously to `lapply()`, by passing an anonymous function. We did not use this approach here, but if we would use this for the MCP home ranges, the call would change from `map(data, hr_mcp)` to `map(data, function(x) hr_mcp(x))`. `x` is just a local variable (i.e., a placeholder) for the current animal’s data and could also be named differently.

The data set `hr1` has now gained a new list column for each home-range estimator (in total there are now five new columns).

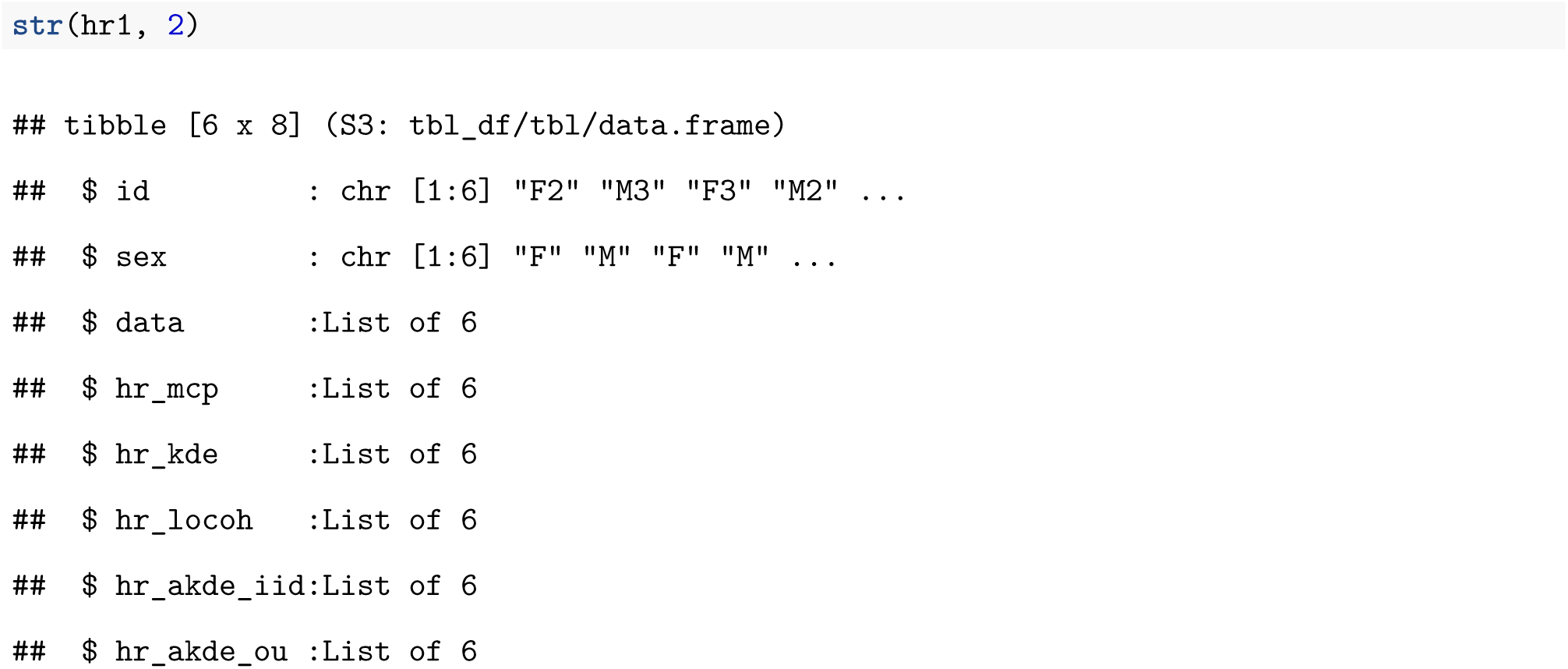

We now want to obtain the home-range size for each animal and each estimator using the same `map`-strategy. Taking advantage of the previously introduced framework, we know that a function `hr_area()` exists for each estimator, and that it will return the home-range size as a `tibble`. However, we would have to apply `hr_area()` separately to each column (`hr_mcp` to `hr_akde_ou`). Instead, we would like to apply the function `hr_area()` to one list containing the home-range estimates for all of the different estimators. To accomplish this task, we need to first change from wide to long format using the function `pivot_longer()` from the package `tidyr` (Wickham and Henry 2020) so that we end up with a tibble that has a column that records the estimator (MCP, KDE, LoCoH, etc) and a second column with the estimates.

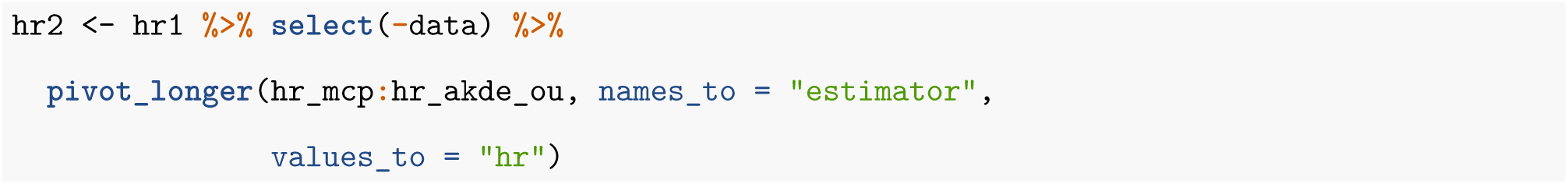

We first removed the tracking data (column `data` as these data are no longer needed) and then pass the resulting `tibble` to the function `pivot_longer`. Here we need to say which columns should be turned from the wide format to the long format (`hr_mcp:hr_akde_ou`). The new data set, `hr2`, will have four columns. The first two columns are `id` and `sex` from the old data set. The third column is called `estimator` (this can be controlled with the argument `names_to`) and identifies the estimator type (i.e., the old column names). The fourth column is called `hr` and contains the actual home-range estimates (the name for this column be controlled again with the argument `values_to`).

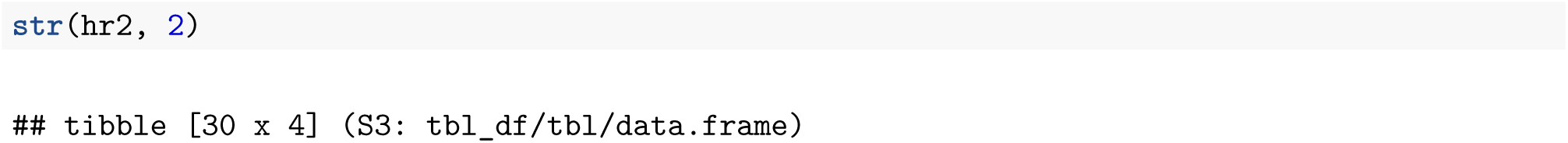

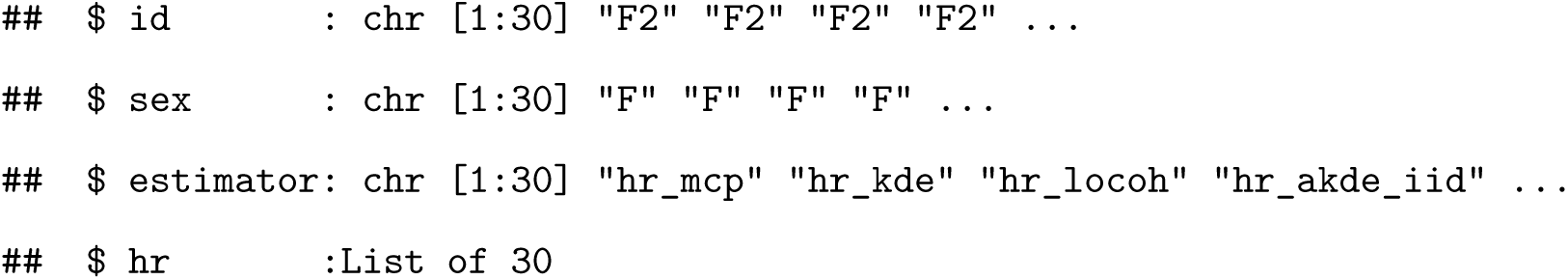

The new long data format allows us to apply the `hr_area` function to each element of the new column `hr` of `hr2`.

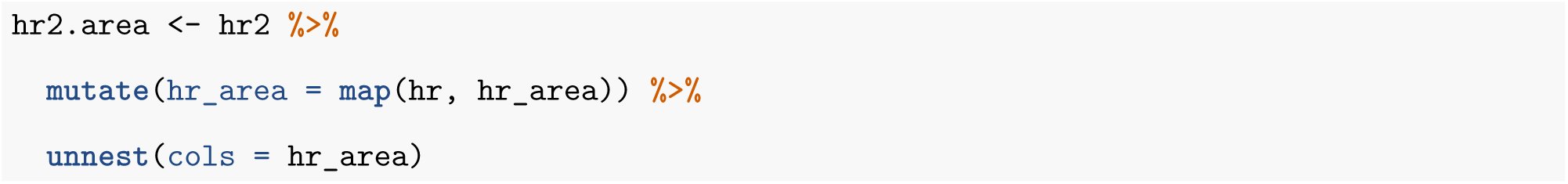

We now undo the list column (with the function `unnest()`). This step is necessary to obtain a `tibble` without a list column, that is suitable for plotting.

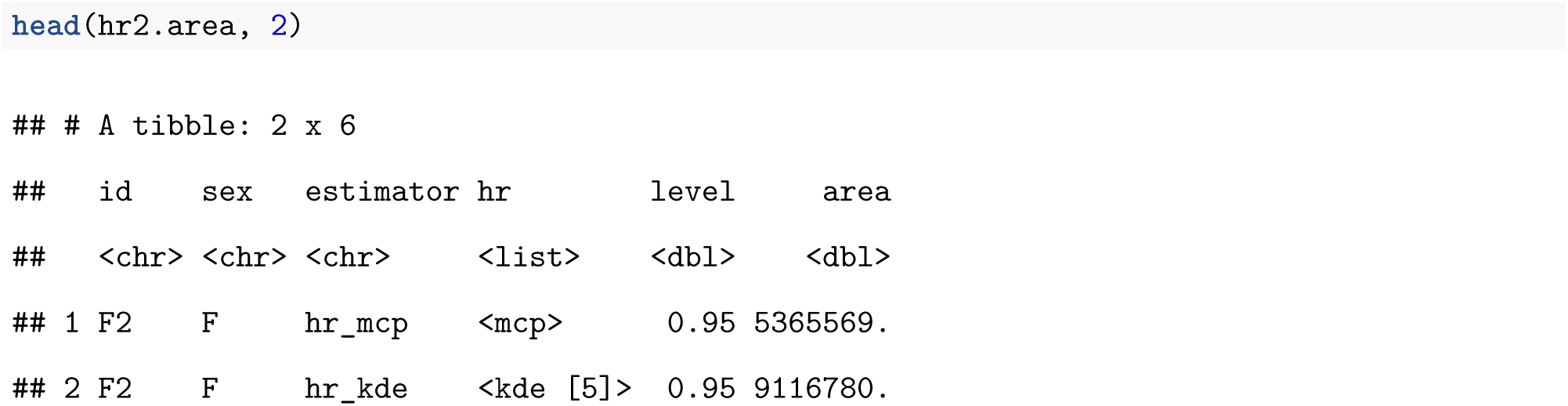

We can now visually explore differences in home-range size between males and females, and consider how these differences are influenced by our choice of home-range estimator (Fig. 3A; full code to reproduce the plot is given in the Supplement 1). Estimates of home-range size differ considerably across the 5 estimators (Fig. 3A). As a result, estimates of the absolute difference in mean home-range size between sexes also varies depending on the chosen estimator (Fig. 3B). By contrast, differences between estimators becomes negligible if we quantify relative differences (i.e., ratios of mean home-range size; Fig. 3C); regardless of estimator choice, we find that male home ranges were 2.4 – 2.8 times larger than female home ranges. With a larger sample of individuals, we could quantify uncertainty in the ratio of mean home-range sizes using a bootstrap (Fieberg, Vitense, and Johnson 2020). If other additional animal-specific covariates were available and of interest, we could use a linear (mixed) model to quantify the relative importance of different covariates in determining home-range size. Again, a larger sample size (i.e., more animals) and a standardized collection scheme would be desirable (Börger, Franconi, De Michele, et al. 2006).

**Figure 3:**
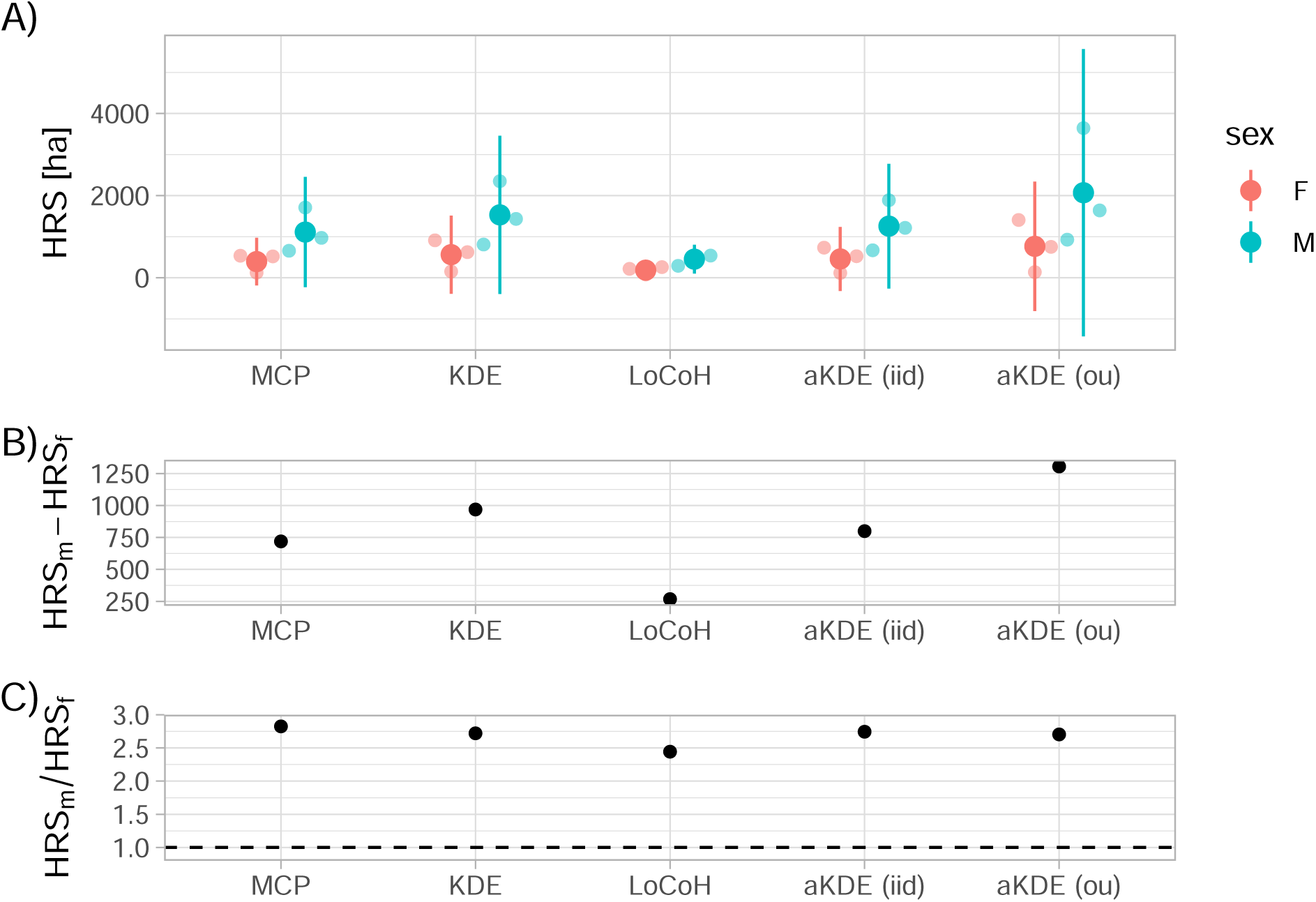
Sexual dimorphism in the size of fisher home-ranges. Different estimators (x-axis) lead to very different home-ranges sizes [HRS] (panel A) and consequently to different estimated absolute differences in HRS between males and females (panel B). However, relative differences (i.e., ratio of mean HRS between males and females) was more consistent, ranging from 2.4 to 2.8 (panel C). In panel A, small dots represent (jittered) individual estimates of HRS, large dots indicate means, and vertical lines represent t-based 95% confidence intervals for each sex. The dashed horizontal line in panel C indicates equivalence of male and female home ranges.

In a second example, we explore whether home-range size correlates with the amount of forest within an animal’s home range. To address this question, we load a preprocessed land use raster (see Supplement 1 for full details) and assign it to the object `env`. As before, we make use of `pivot_longer` to obtain one list column with all home-range estimates and then obtain the isopleth levels with `hr_isopleth()` function. Again this works for all implemented estimators in the package `amt` and results in an `sf`-object. With the function `extract` from the raster package (Hijmans 2020), the pixel values within each of the home-range isopleths can be queried.

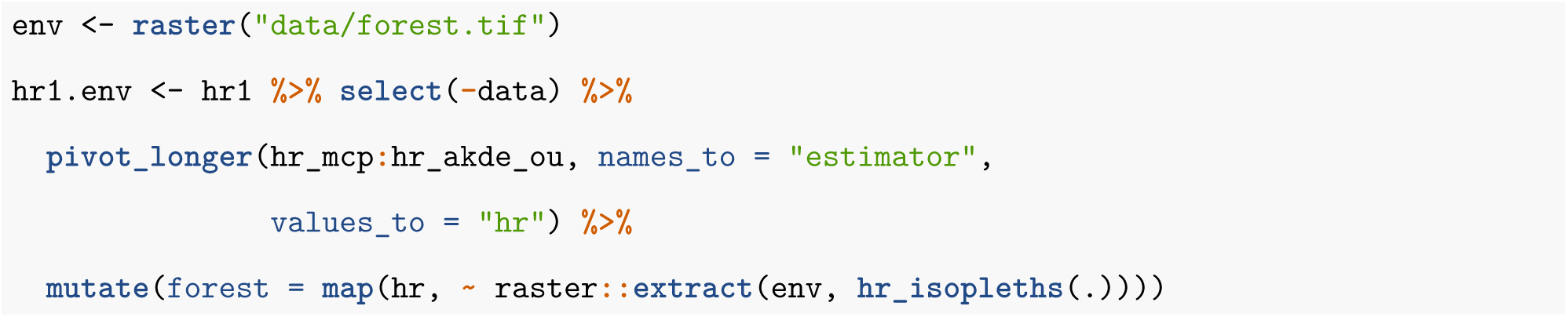

In a final step for this analysis, we calculate the proportion of forest pixels within each individual’s estimated home range and the home-range size. For the proportion, we iterate again over the list `env`, but this time we use the function `map_dbl()`, a variant of `map()` that will always return a numeric vector.

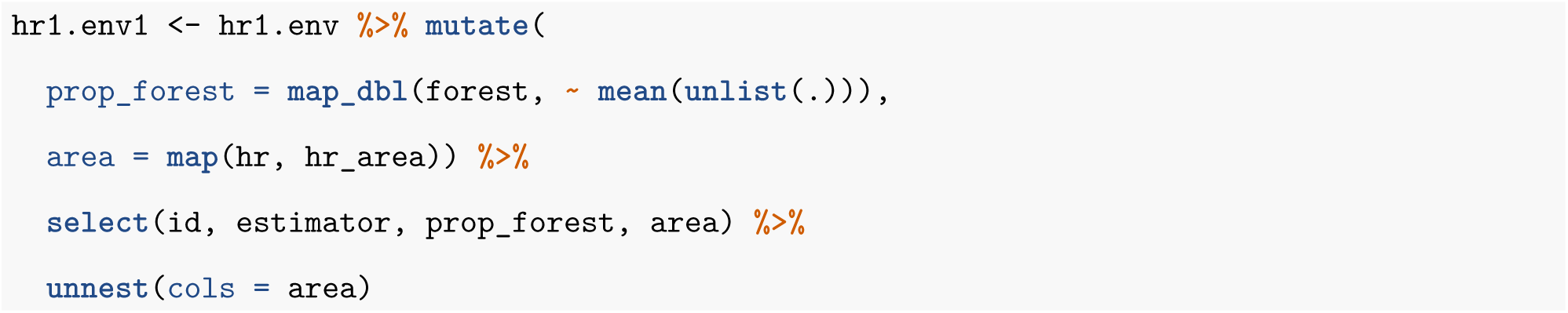

We need to make a call to the function `unlist()` within the function `mean()` because `extract()` returns a list, allowing for more than one polygon per feature as is common with some home-range estimators (e.g., LoCoH). For this application, however, we can safely combine the land cover classes for different polygons belonging to the same animal. We use the resulting `tibble hr1.env1` to plot estimates of home-range size against the proportion of the estimated home-range composed of forest (Fig. 4). Similar to the previous example (Fig. 3), we observe that different home-range methods result in vastly different estimates in absolute terms, but the observed pattern (i.e., home-ranges with more forest tend to be larger in size) is consistent among all estimators. In situations with more tracked animals and more (environmental) covariates, linear (mixed) models could be used to simultaneously explore multiple determinants of home-range size.

**Figure 4:**
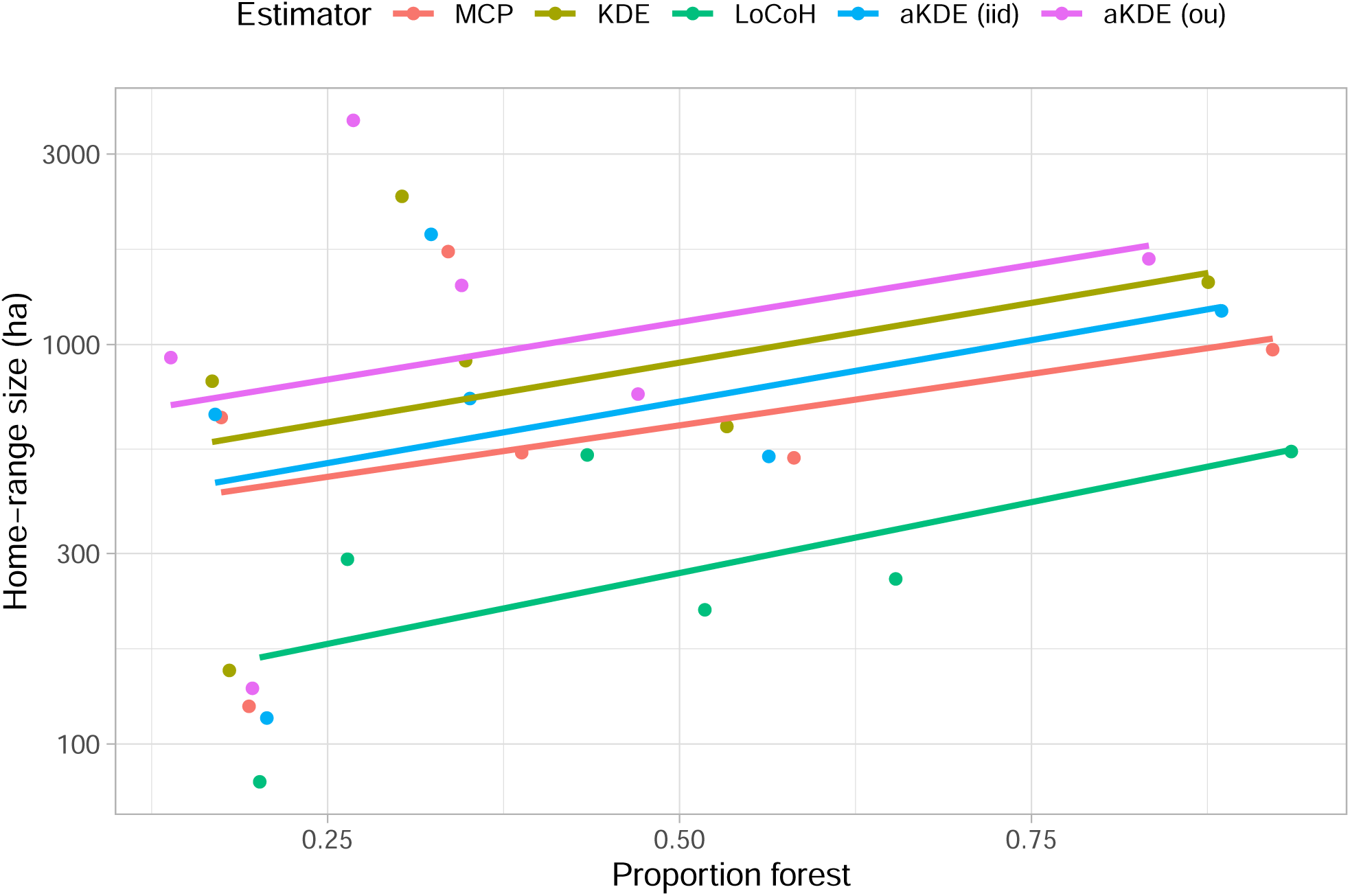
Changes of home-range size as a function of proportion of forest within the home range with linear trend lines. Different estimators (line colors) lead to very different absolute home-range sizes (y-intercepts) but very similar trends (slope of the lines).

Lastly, we consider an example exploring the extent to which individual space-use patterns change over time. To do so, we first add a new column to the `tibble` with the day of the year (called `yday`) and then group our data set by animal id (`id`) and the day of the year (`yday`). This results in a new `tibble` where the relocations for each animal and year-day are stored in the column `data`. We will only consider days with > 10 relocations. To do this, we first count the number of relocations of each instance (animal-year-day) and then filter for days with > 10 relocations. We then calculate five different home-range estimates as before, but for the new grouping and swapping out Fleming et al. (2015)’s aKDE estimator of the range distribution with Fleming et al. (2016)’s estimator of the occurrence distribution [OD]; the latter is more appropriate for use with short tracking periods where interest lies in estimating the actual path of the animal rather than its equilibrium distribution.

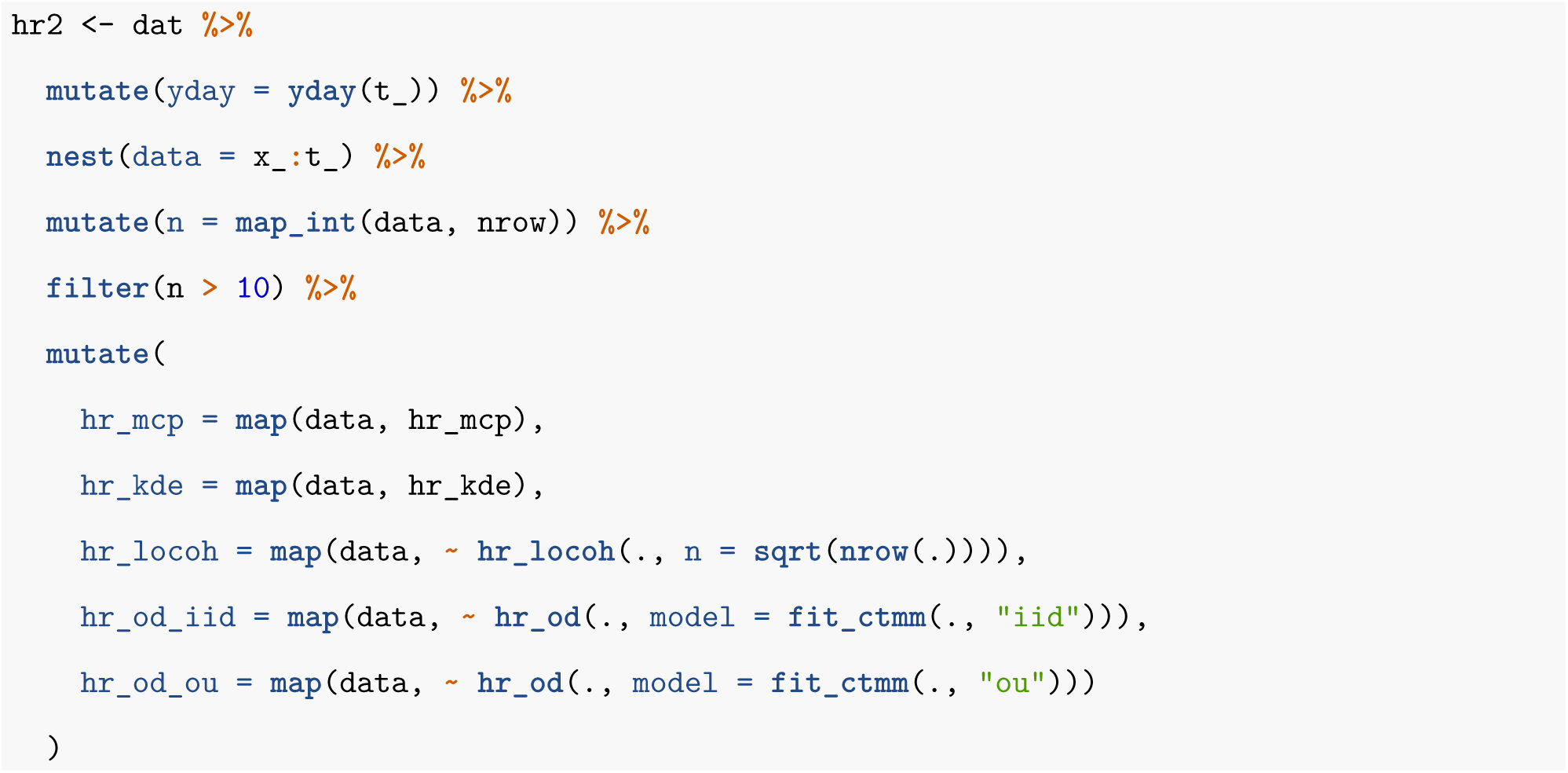

We then follow the same design pattern as before, combining all animals into one list column in the long format and applying the function `hr_area` to all estimates. As with the other 2 examples, there are large consistent differences between the 5 estimators, but all exhibit similar trends over time (Fig. 5). The full code is again given in the Supplement 1.

**Figure 5:**
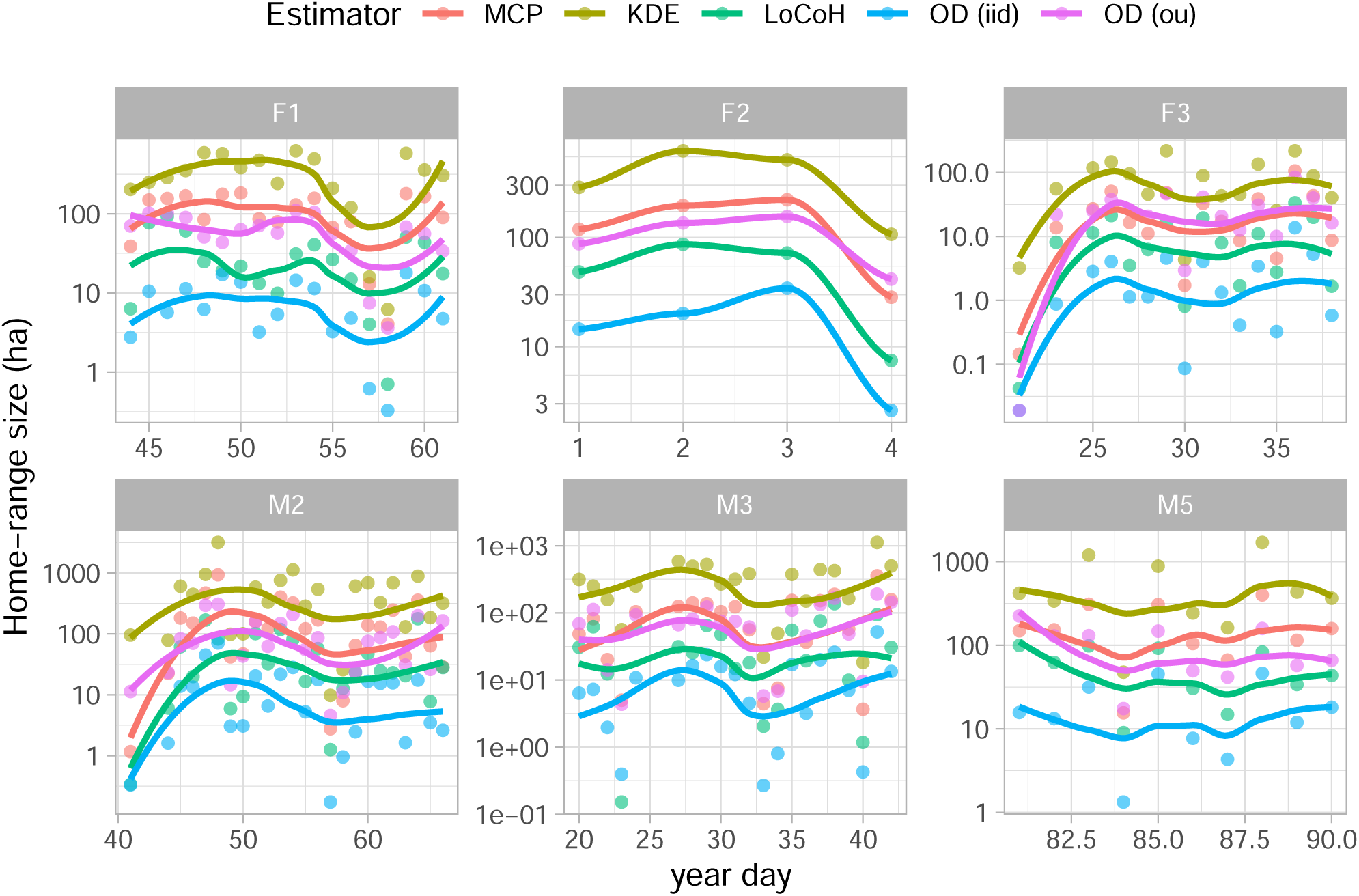
Daily space-use indices for six different fisher estimated using 5 different estimators, along with smooth temporal trends estimated with `ggplots geom_smooth()` function.

## Discussion

The home range is an important biological concept that has and will continue to be highly influential. Fieberg and Börger (2012) argued we should clearly distinguish the biological concept of a home range from the statistical methods used to gain insights into this concept. It is also important to recognize that home-range estimates are not always the end goal, but rather, estimates of home-range size are often used to explore questions regarding how various factors influence animals’ use of space (Börger, Franconi, De Michele, et al. 2006; Börger, Franconi, Ferretti, et al. 2006b); often, these questions involve comparisons of home-range estimates over space or time and for different population segments. For example, researchers may correlate estimates of home-range size with demographic traits, landscape features that also vary in time, or the presence or absence of predator species (Beest et al. 2011; Tingley et al. 2014; Ditmer et al. 2018). Estimates of animal home-ranges are also often used to determine habitat availability when studying habitat selection. Although we agree with Fleming et al. (2016) and Horne et al. (2020) that *range* and *occurrence* distributions are useful estimation targets that can help end users choose an appropriate statistical home-range method, for some situations it may not be clear which of the two concepts (if either) is most suitable for addressing a particular research question. For example, a range distribution will not be appropriate for studying temporarily varying space-use patterns. On the other hand, researchers may want to incorporate areas that were likely known and accessible to the animal but not used during a specific observation window when studying habitat selection (i.e., they may be interested in more than just the animal’s movement path, which would be estimated by the occurrence distribution).

Analysts often seek simple measures to detect changes in spatial extent of movements over time, between sexes or habitats. Like Signer et al. (2015), we found that answers to questions that involve relative comparisons of home-range size were robust to estimator choice. Yet, differences in the tracking regime (VHF or GPS) and sampling rate (i.e., how often is an animal tracked) can lead to vastly different home-range estimates depending on one’s choice of estimator (Noonan et al. 2019; Peris et al. 2020). These differences can also influence estimates of derived quantities and observed relationships, for example scaling laws between home-range size and body size (e.g., Noonan et al. 2020). Thus, it is important for researchers to conduct sensitivity analyses to determine how choice of estimator influences their quantitative and qualitative results. The standards we outline here should make this task simple to accomplish.

In the spirit of allowing users to freely apply and evaluate multiple estimators, we suggest approaching home-range estimation in a consistent manner, using a coherent and tidy work flow that facilitates quantification of space use of different animals when various grouping instances are present (e.g., individuals, temporal units such as days or weeks, or both). Animal tracking is still in its early stages (Kays et al. 2015), standards for tracking data are still in flux (Campbell et al. 2016), and new methods are constantly being developed. As wildlife biology enters an era of big data, a coherent, scriptable and reproducible workflow is needed to ensure reproducibility of results (Lewis, Vander Wal, and Fifield 2018; Archmiller et al. 2020), and a standardized, tidy implementation of home-range estimators should help facilitate that vision.

## Acknowledgments

J.Fieberg received partial salary support from the Minnesota Agricultural Experimental Station.

## Authors contribution

JS and JF conceived the ideas and designed methodology; JS analyzed the data; JS and JF led the writing of the manuscript. Both authors gave final approval for publication.

## Data availability

All data used in this manuscript were already published an are available here: https://zenodo.org/record/3991482.

